# Characterising retinal function with optomotor visual performance in P23H rodent models of retinitis pigmentosa

**DOI:** 10.64898/2026.04.09.717562

**Authors:** Alicia A. Brunet, Daniel Urrutia-Cabrera, Luozixian Wang, Genevieve Huppert, Stephanie Chu, Rebekah James, Alan R. Harvey, Raymond C.B. Wong, Livia S. Carvalho

**Affiliations:** Lions Eye Institute Ltd., 2 Verdun St, Nedlands, WA 6009, Australia; Centre for Ophthalmology and Visual Sciences, The University of Western Australia, 35 Stirling Hwy, Crawley, WA 6009, Australia; Department of Optometry and Vision Sciences, University of Melbourne, Parkville 3052, Victoria, Australia; Centre for Eye Research Australia, Royal Victorian Eye and Ear Hospital, Australia; Ophthalmology, Department of Surgery, University of Melbourne, Australia; School of Human Sciences, The University of Western Australia, 35 Stirling Hwy, Crawley, WA 6009, Australia; Perron Institute for Neurological and Translational Science, 8 Verdun St, Nedlands, WA 6009, Australia

**Keywords:** optomotor, retinitis pigmentosa, inherited retinal diseases, photoreceptor degeneration, rodent disease models

## Abstract

Rhodopsin (RHO) P23H is one of the most common mutations causing autosomal dominant retinitis pigmentosa (adRP), yet the relationship between retinal electrophysiology, structure and visually guided behaviour in rodent models remains unclear. We characterised changes in heterozygous P23H (Sakami line) mice and P23H line 3 (P23H-3) rats using full-field electroretinography (ERG), optomotor response (OMR) assays and, in rats, optical coherence tomography (OCT). ERG assessed rod- and cone-mediated responses relative to wild-type controls, whereas OMR under scotopic and photopic conditions quantified contrast sensitivity and visual acuity. In P23H mice, scotopic ERG responses were significantly reduced from postnatal day 16 and declined further from 4 months. Scotopic OMR contrast sensitivity remained largely preserved until 2 months, and photopic acuity was comparable to wild-type up to 6 months. In 13-week-old P23H-3 rats, ERG amplitudes were significantly reduced, and OCT revealed retinal thinning. OMR showed a decline in contrast sensitivity at 7 and 15 weeks, whereas photopic acuity was maintained. Thus, in both models, electrophysiological and structural abnormalities precede detectable OMR deficits, with implications for the selection of outcome measures in preclinical studies.

**Summary Statement:** This study compares electrical and behavioural measures of vision in rodent models of inherited blindness, revealing that retinal dysfunction appears well before measurable vision loss.

## Introduction

Retinitis pigmentosa (RP) is the most common form of inherited retinal disease (IRD), affecting 1 in 4,000 individuals worldwide (Wang et al., 2019). RP is a genetically heterogenous condition, with mutations in over 100 genes identified to cause the disease (Rivolta et al., 2025). These mutations primarily arise in rod photoreceptors, resulting in rod degeneration, followed by a progressive mutation-independent loss of cone photoreceptors. As such, patients initially present with night blindness. As the disease advances, a progressive deterioration of peripheral daylight vision and visual acuity also occurs (Verbakel et al., 2018).

A point mutation in the rhodopsin gene (*RHO*) that results in a proline-to-histidine substitution at codon 23 (P23H) is one of the major mutations associated with autosomal dominant RP (Dryja et al., 1990). This mutation causes the misfolding of rhodopsin proteins, which can trigger endoplasmic reticulum stress and the unfolded protein response leading to rod degeneration (Yao et al., 2018, Liu et al., 2020, Gorbatyuk et al., 2010). Mouse and rat models with the P23H rhodopsin mutation recapitulate many of the pathophysiological and molecular changes observed in patients with autosomal dominant RP (Barwick and Smith, 2023, LaVail et al., 2018). Both the homozygous (*Rho^P23H/P23H^*) and heterozygous (*Rho^P23H/WT^*) forms have been investigated in various animal models such as mice (Sakami et al., 2011, Price et al., 2011), rats (LaVail et al., 2018, Orhan et al., 2015, McGill et al., 2012) and pigs (Ross et al., 2012). There are four variations of P23H mouse model generated by different labs, though the Sakami *et al*. (2011) model (available from The Jackson Laboratory #017628) is thought to mimic the human disease better than other models (Sakami et al., 2011, Sakami et al., 2014, Barwick and Smith, 2023). *Rho^P23H/P23H^* mice have been shown to have a severe phenotype, with rapid photoreceptor degeneration and loss of visual function (Sakami et al., 2011, Sakami et al., 2014). On the other hand, *Rho^P23H/WT^* have demonstrated a more gradual degeneration, more closely representing the human inheritance pattern and phenotype. Similarly, the ratio of mutant to wild type rhodopsin expression is critical in determining the degeneration rate in transgenic *Rho^P23H/WT^*rats. In 2018, LaVail *et al*. reported the production of three transgenic rat lines with the introduction of a mouse rhodopsin gene with the P23H mutation (LaVail et al., 2018). Line 1 (P23H-1) expressed the highest levels of P23H mutated rhodopsin, resulting in the earliest onset (P10) and fastest photoreceptor degeneration rate from the three lines. In contrast, line 3 (P23H-3) presented a moderate rate of photoreceptor degeneration, mimicking the human disease progression. Therefore, selecting P23H animal models that most closely align with the clinical findings of the human P23H presentation is critical for disease modelling as well as therapeutic development.

The optomotor response (OMR) is a non-invasive behavioural assessment of visual function (Kretschmer et al., 2017). Unlike electroretinography (ERG), which measures retinal electrical activity, OMR integrates the whole visual pathway, providing essential information about the effect of photoreceptor degeneration in vision loss. OMR testing in *Rho^P23H/WT^* mice has given valuable insights into visual behaviour and degeneration associated with this form of RP. Interestingly, OMR testing in 1.5-month-old *Rho^P23H/WT^* mice (Sakami line) revealed ‘supernormal’ cone-mediated visual behaviour compared to wildtype at low spatial frequencies, as well as hyperexcitability to visual stimuli at the primary visual cortex level (Leinonen et al., 2022). This trend was maintained up to 3 months of age, albeit with slight reductions in OMR frequency, aligning with the progressive degeneration observed in this model. Another study demonstrated no change in contrast sensitivity in *Rho^P23H/WT^* mice and suggested that compensatory mechanisms within the retina may help to maintain night vision, despite photoreceptor degeneration, up to 5 months of age (Leinonen et al., 2020). However, they do note that this compensatory effect diminishes with disease progression. McGill *et al*. (2012) also performed OMR testing in P23H line 1, 2 and 3 pigmented rats (McGill et al., 2012). Interestingly, similar to some previous mouse studies, this decline in visual function was observed much later in the degeneration timeline, after significant photoreceptor loss had already occurred. The authors suggest that for future behavioural studies, prior to therapeutic intervention, careful analysis of the visual thresholds of animals is needed. Additionally, these findings in different P23H rats may also allude to potential compensatory mechanisms within the retina, particularly in the earlier stages of the disease.

The present study aims to characterise and provide a direct comparison of the *Rho^P23H/WT^* mouse (Sakami line) and rat (P23H-3) models of autosomal dominant RP that most closely recapitulate the human disease phenotype, directly comparing ERG data with the more global assessment of visual function using OMR. OMR systems with varying designs and mode of operation have previously been used to study OMR in rodents, which may yield variations in OMR measurements. As such, we utilised the automated OptoDrum (Striatech, Germany) system to eliminate user-dependent scoring biases associated with manual OMR setups. We show that ERG responses in *Rho^P23H/WT^*rodents were significantly decreased at all ages tested, though decreases in OMR contrast sensitivity were only evident after substantial rod degeneration had reportedly occurred. Our findings highlight the sensitivity of OMR to detect residual visual function, even at stages when ERG responses are diminished or absent.

## Results

### ERG response changes in Rho^P23H/WT^ mice over time

To establish the extent of functional degeneration in P23H rodent models, full-field ERG was performed as an assessment of retinal electrophysiological activity. ERGs were conducted on *Rho^P23H/WT^* mice to evaluate age-related changes in retinal response over time compared to healthy wildtype controls. Scotopic responses at the highest light intensity (25 cd.s/m^2^) are shown in Figure 1 (all light intensities shown in Supplementary Figure 1). Scotopic ERG responses matured around 1 month of age in wildtype mice, consistent with previously reported developmental timelines (Gibson et al., 2013), though *Rho^P23H/WT^* mice may have a slightly later maturation at P40. Responses then decreased with age in both lines, with previous literature establishing that ERGs begin decreasing as early as 5 weeks of age in wildtype C57BL/6 mice (Park et al., 2023). However, *Rho^P23H/WT^* mice had a significantly reduced ERG response at all timepoints. Between P16 to 2 months *Rho^P23H/WT^* mice had a consistent 50-60% reduction in scotopic a-wave compared to age-matched wildtype (Figure 1, Supplementary Figure 2). By 4 months of age, this reduction was even greater at 78%, suggesting ongoing degenerative changes occurring between 2 and 4 months of age. There was also a greater reduction in scotopic b-waves in *Rho^P23H/WT^* mice at 4 months with a 64% reduction. To benchmark the sensitivity of ERGs in detecting retinal responses during advanced degeneration, we also evaluated the autosomal recessive *rd1* mouse model, which exhibits rapid photoreceptor loss (Keeler, 1924). Both a-wave and b-wave scotopic ERG responses in *rd1* mice were severely reduced at postnatal day 16 (P16) compared to wildtype mice and no longer detectable by P24 (Figure 1). Oscillatory potentials (OPs) were also evaluated as a further assessment of the disturbances to inner retinal function during disease and showed decreases in *Rho^P23H/WT^*mice compared to wildtype (Supplementary Figure 3). Representative waveforms of OPs depict a latency in *Rho^P23H/WT^* OPs.

**Figure 1.**
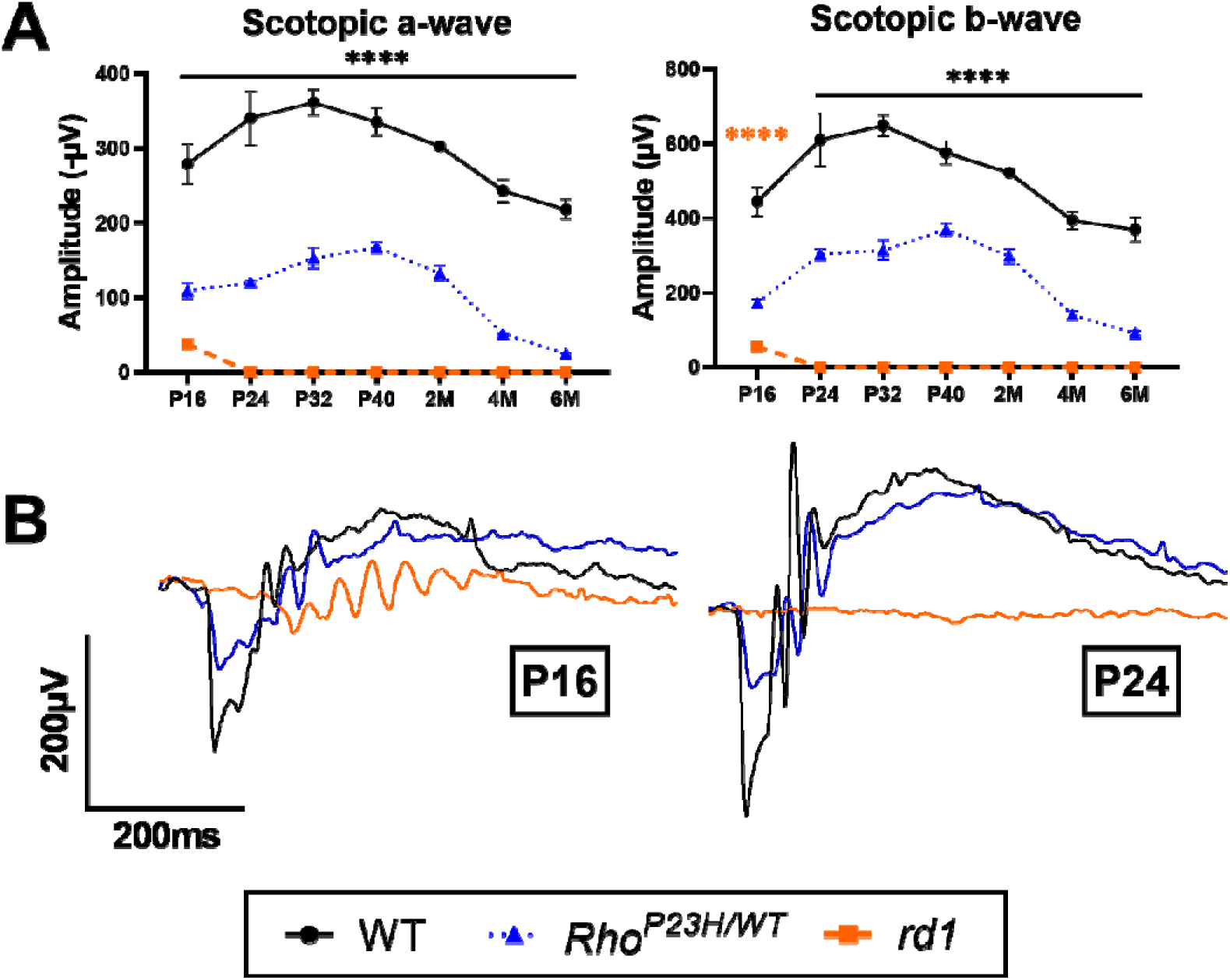
Scotopic ERG responses in *Rho^P23H/WT^* and *rd1* mice compared to wildtype at 25 cd.s/m^2^. (**A**) Scotopic a- and b-waves in *Rho^P23H/WT^*and *rd1* mice compared to wildtype mice across age. Black line denotes both *Rho^P23H/WT^* and *rd1* mice were significantly decreased compared to wildtype. Orange asterisks denotes a significant decrease in *rd1* mice compared to wildtype. Numbers are mean ± SEM. *n* = 3-6 per age per line. \*\*\*\**P* < 0.0001, two-way ANOVA and Tukey’s multiple comparisons test. 2M: 2 months; 4M: 4 months; 6M: 6 months. (**B**) Representative traces of scotopic ERG responses for each line at P16 and P24.

To assess cone-mediated visual function during degeneration, photopic ERG responses were recorded in *Rho^P23H/WT^* and *rd1* mice compared to wildtype controls. Figure 2 shows responses at 25 cd.s/m^2^, and all light intensities are shown in Supplementary Figure 4. *Rho^P23H/WT^* mice had comparable photopic a-wave responses to wildtype mice at all timepoints except P24, although there was a trend for decreased responses. The photopic b-wave was significantly decreased at every timepoint except P16 and 2 months of age. In *rd1* mice, the photopic a-wave was comparable to wildtype mice at P16 whilst the photopic b-wave was significantly decreased, but both were diminished to an unmeasurable level by P24. The complete loss of photopic responses by P24 mirrors the loss of scotopic responses at this age. Representative photopic waveforms are provided for P16 and P24 to show the loss of response at P24. Photopic waveforms also showed an observable latency in response in *rd1* mice (indicated by black arrow).

**Figure 2.**
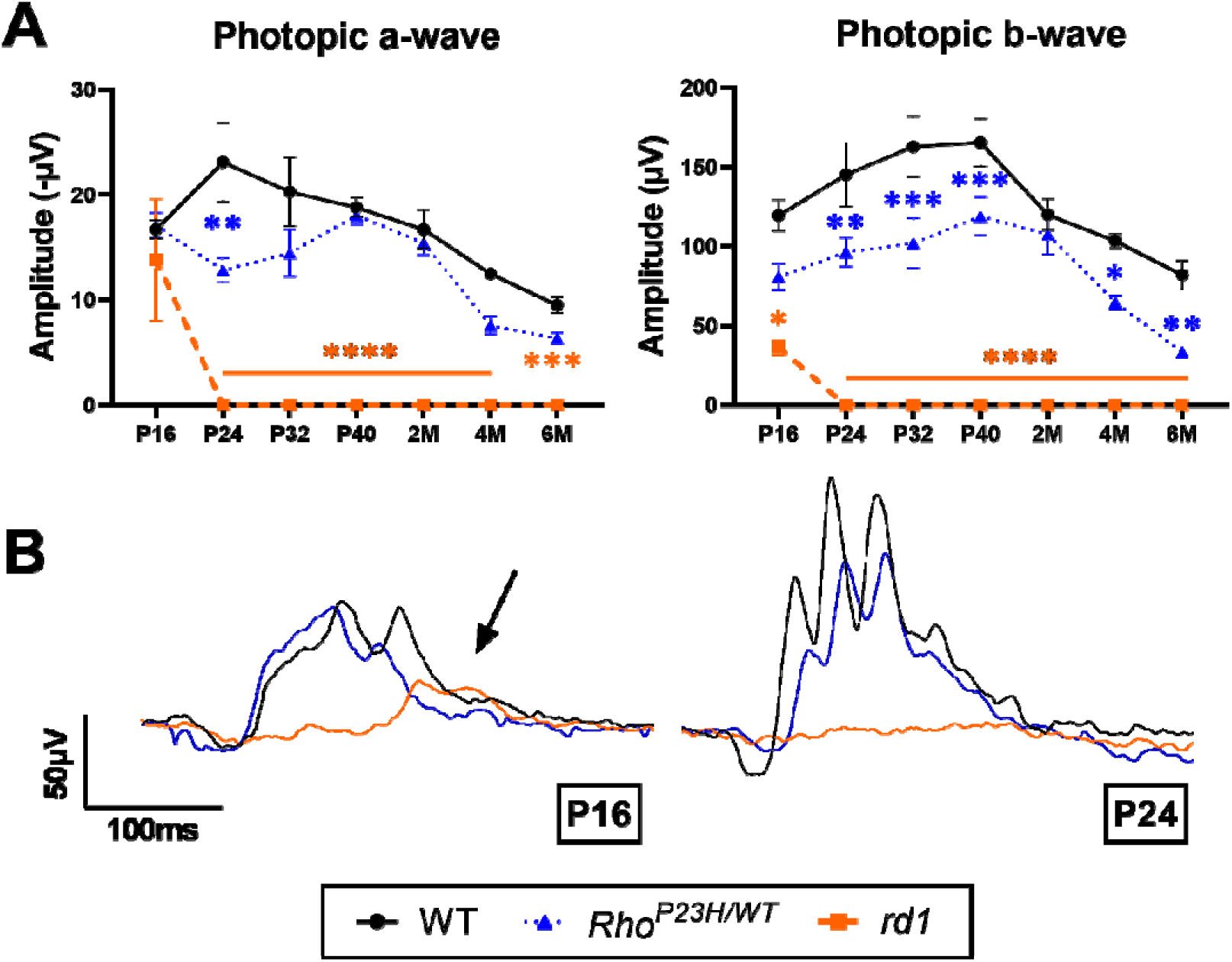
Photopic ERG responses in *Rho^P23H/WT^* and *rd1* mice compared to wildtype at 25 cd.s/m^2^. (**A**) Photopic a- and b-waves in *Rho^P23H/WT^* and *rd1* mice compared to wildtype. Significant decreases in *Rho^P23H/WT^*and *rd1* mice compared to wildtype are shown by blue and orange asterisks, respectively. Numbers are mean ± SEM. *n* = 3-6 per age per line. \**P* < 0.05, \*\**P* < 0.01, \*\*\**P* < 0.001, \*\*\*\**P* < 0.0001, two-way ANOVA and Tukey’s multiple comparisons test. 2M: 2 months; 4M: 4 months; 6M: 6 months. (B) Representative traces of photopic responses are shown for P16 and P24. The *rd1* mouse had a notable delay in photopic response (indicated by black arrow).

### Optomotor response analysis in Rho^P23H/WT^ mice with visual impairment

The OMR is a compensatory head reflex evoked by a moving stimulus and is a metric to assess visual performance. Despite significantly reduced rod-mediated ERG responses, contrast sensitivity in *Rho^P23H/WT^*mice under scotopic conditions was only significantly decreased compared to wildtype mice at 2 months and 4 months of age. Additionally, although not statistically significant, there was also a trend towards reduced OMR response at 6 months of age (Figure 3B). This significant decrease at 2 months coincides with the previously reported timepoint whereby half the population of rods have degenerated in *Rho^P23H/WT^*mice (Sakami et al., 2011). Visual acuity under photopic conditions in *Rho^P23H/WT^*mice was similar to wildtype mice at all ages measured. In comparison, *rd1* mice were unable to detect the highest contrast setting (99.97%) in scotopic conditions from P16 onwards (data not shown). This lack of rod-mediated optomotor response reflects the severely decreased scotopic ERG response in *rd1* mice. In photopic conditions, *rd1* mice were able to detect the stimulus at P16 and P24, albeit significantly decreased from wildtype mice. *Rd1* mice were therefore able to still respond to moving stimuli at P24 despite the lack of ERG response at this age, indicating a higher sensitivity of the OMR to residual photoreceptor activity compared to full-field ERGs. By P32, *rd1* mice no longer had an optomotor response even to the highest acuity setting (0.0556 cyc/deg).

**Figure 3.**
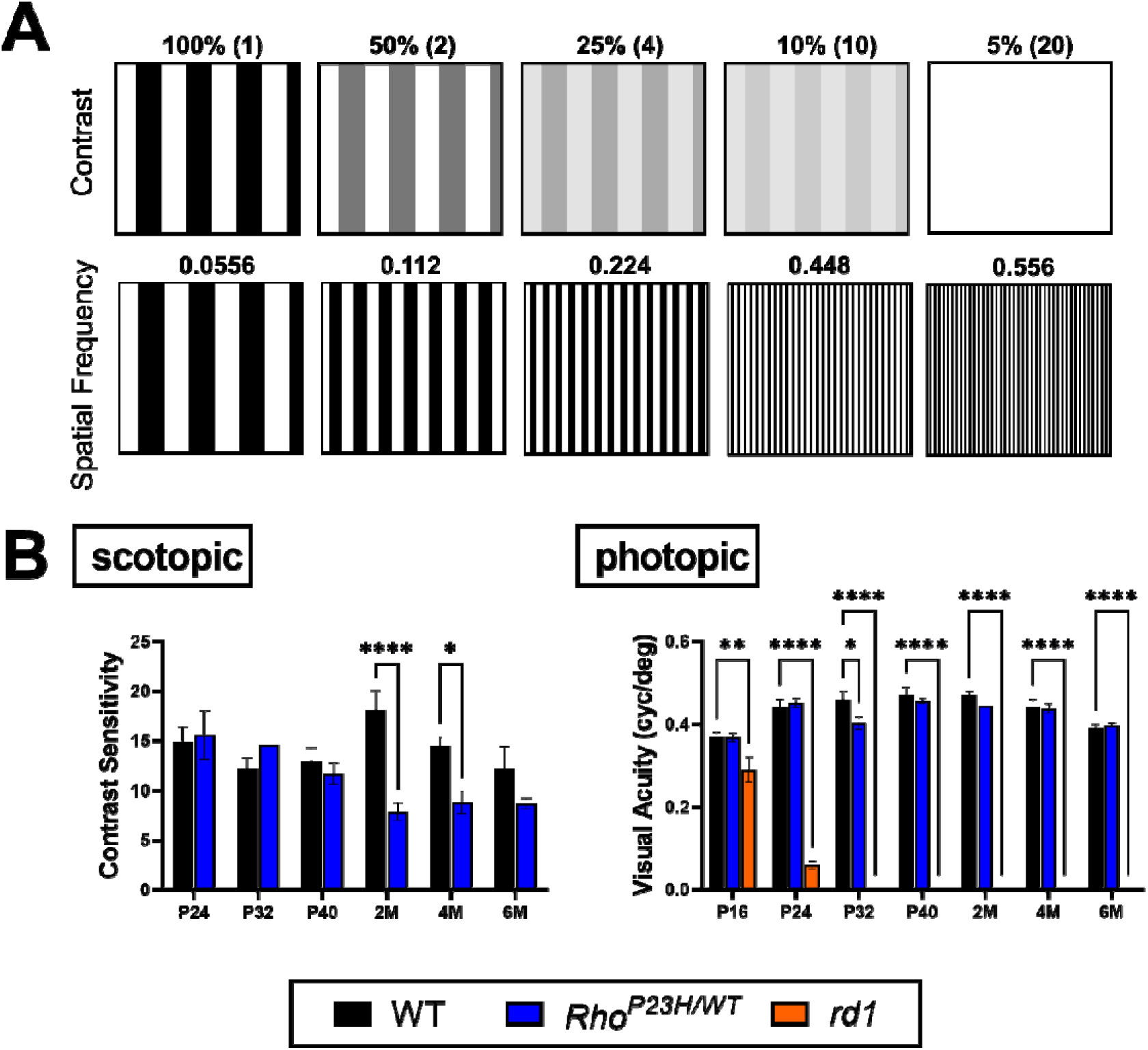
Optomotor responses of *Rho^P23H/WT^* and *rd1* mice compared to wildtype across ages. (**A**) Examples of different contrasts (presented as inverse of contrast percentage) and spatial frequencies (measured as cycles per degree, cyc/deg) in the optomotor tests. (**B**) Optomotor contrast sensitivity in scotopic conditions and visual acuity in photopic conditions of all lines. Numbers are mean ± SEM. *N* = 4-9, per age per line. \**P* < 0.05, \*\**P* < 0.01, \*\*\*\**P* < 0.0001, two-way ANOVA and Tukey’s multiple comparisons test. 2M: 2 months; 4M: 4 months; 6M: 6 months.

### Characterisation of photoreceptor degeneration in the transgenic P23H-3 rat

The transgenic P23H-3 rat is characterised by a slow to moderate rate of photoreceptor degeneration with changes in the ONL thickness reported to be distinguishable at 6 months of age in pigmented rats (LaVail et al., 2018). Notably, since albino rats showed poor optomotor reflex in our initial tests (Supplementary Figure 5A), we crossed our albino P23H-3 rats in Sprague-Dawley (SD) background with pigmented Long-Evans rats. Since there was a marked reduction in scotopic ERG responses between 2 and 4 months of age in *Rho^P23H/WT^* mice, we chose the approximate midpoint of this interval (13 weeks) as a key timepoint for assessment in P23H-3 rats. Scotopic ERG analysis determined that at 13 weeks of age, P23H-3 rats showed a significant decrease in a-wave amplitude compared to wildtype (SD) rats at all the light intensities tested (Figure 4A). These results indicate a reduction primarily of rod photoreceptor response, which is consistent with the pathophysiology observed in RP patients. Similarly, the scotopic b-wave in P23H-3 rats was significantly reduced compared to wildtype rats (Figure 4B). Interestingly, the ERG response was reduced at higher rate for low intensity flashes including both a-wave and b-wave. In all light intensities tested, photoreceptor function (a-wave) was more affected in P23H-3 rats with a 63-78% reduction (Supplementary Figure 6A), compared to a 37-48% decrease in b-wave response (Supplementary Figure 6B). Overall, the traces of scotopic ERG response were reduced in P23H-3 rats compared to wildtype even in response to the highest flash intensity tested (Figure 4C).

**Figure 4.**
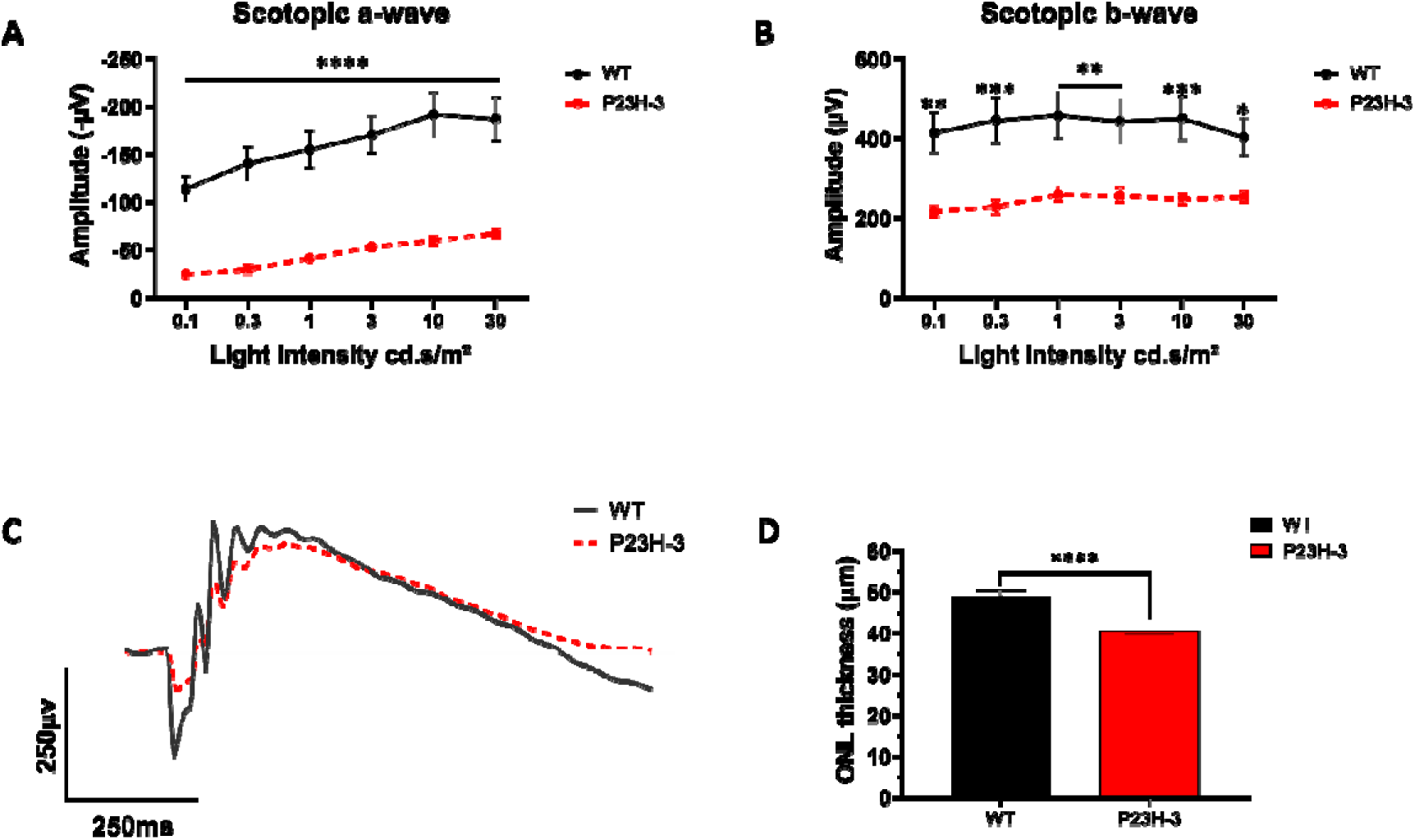
Analysis of retinal function and thickness in P23H-3 rats. Scotopic ERG analysis showing the (**A**) a-waves and **B**) b-waves detected in wildtype and P23H-3 rats in response to different light intensities. At 13 weeks of age, P23H-3 rats (red) showed a significant decrease in both scotopic a-wave and b-waves compared to wildtype rats (black). Numbers are mean ± SEM. n = 10-12 eyes. \**P* < 0.05, \*\**P* < 0.01, \*\*\**P* < 0.001, \*\*\*\**P* < 0.0001, one-way ANOVA. **C**) Representative traces of scotopic responses at 13 weeks of age for wildtype (black) and P23H-3 rats (red). Lines represent the mean of the response for *n* = 10-12 eyes. (**D**) OCT analysis comparing the ONL thickness in P23H-3 and wildtype rats at 13 weeks of age. Numbers are mean ± SEM. *n* = 10-12 eyes, per age per line. \*\*\*\**P* < 0.0001, unpaired t-test.

A decrease in retinal function likely indicates changes in retina structure triggered by the dysfunction and degeneration of rod photoreceptors. Indeed, a reduction in outer nuclear layer (ONL) thickness has been extensively reported in P23H rodent models, though the rate of photoreceptor degeneration differs between albino and pigmented strains (LaVail et al., 2018, McGill et al., 2012). Consistent with previous reports, we identified a significant reduction of ONL thickness in pigmented P23H-3 rats at 13 weeks of age compared to wildtype (Figure 4D). The ONL thickness in wildtype rats measured with OCT was 49.04 ± 1.25 µm, while P23H-3 rats had a significantly thinner ONL of 40.56 ± 1.08 µm. These results indicate that P23H-3 rats showed impaired retinal structure and function at the age of 13 weeks, demonstrated by a greater reduction of the photoreceptor ERG response and a decrease of the ONL thickness.

### OMR analysis in P23H-3 rats demonstrate visual impairment

Having shown that P23H-3 rats at 13 weeks of age experience a decrease in ERG responses, we further investigated whether these changes could also affect behaviours associated with visual function such as the OMR. Similarly to the *Rho^P23H/WT^* mice, we studied changes in contrast sensitivity (Figure 5A) and visual acuity (Figure 5B). We first performed a contrast sensitivity curve in scotopic conditions to study the effect of spatial frequency in the contrast sensitivity of wildtype and P23H-3 rats at 15 weeks of age (Figure 5C). Both groups showed a maximum contrast sensitivity at a spatial frequency of 0.083 cyc/deg, which was reduced at higher values of spatial frequency. In general, the contrast sensitivity of P23H-3 rats was slightly lower at every spatial frequency compared to wildtype.

**Figure 5.**
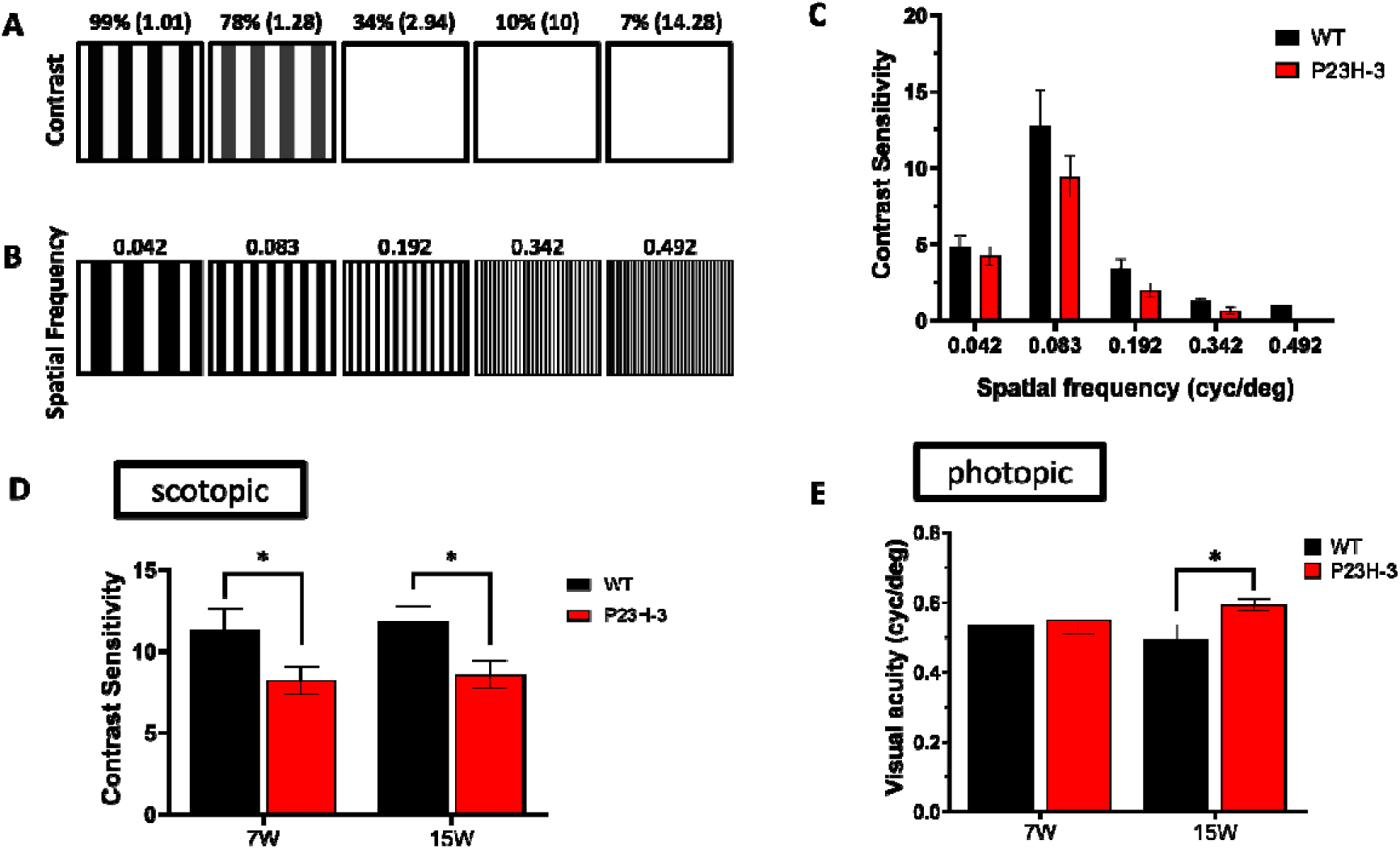
Optomotor response of P23H-3 rats compared to wildtype. Schematic representation of different (**A**) contrast percentages and (**B**) spatial frequencies (cycles per degree, cyc/deg) presented to the rats during the optomotor reflex tests. (**C**) Contrast sensitivity curve as a function of the spatial frequency in P23H-3 and WT rats tested under scotopic conditions. (**D**) Scotopic contrast sensitivity analysis in P23H-3 rats at 7 and 15 weeks of age, showing a significant decrease compared to WT rats. Numbers are mean ± SEM. *n* = 6-19 eyes (WT) and *n* = 12-22 eyes (P23H-3). \**P* < 0.05, unpaired t-test. (**E**) Photopic visual acuity test in P23H-3 rats at 7 and 15 weeks of age compared to WT. Numbers are mean ± SEM. *n* = 6-22 eyes (WT) and *n* = 10-22 eyes (P23H-3). \**P* < 0.05, unpaired t-test. 7W: 7 weeks; 15W: 15 weeks.

Scotopic OMR analysis identified a decrease in contrast sensitivity at 7 weeks of age with P23H-3 rats showing 8.21 ± 0.85 and wildtype rats presenting 11.31 ± 1.31 of contrast sensitivity. Notably, the contrast sensitivity determined at 15 weeks remained largely unchanged for both rat strains, though the contrast sensitivity of P23H-3 (8.55 ± 0.82) was still significantly lower compared to wildtype rats (11.83 ± 0.91) (Figure 5D). In contrast, the visual acuity of P23H-3 rats was comparable to wild type rats at 7 weeks of age (Figure 5E). Our results showed that the visual acuity of P23H-3 rats ranged from 0.55 ± 0.04 cyc/deg at week 7 to 0.59 ± 0.01 cyc/deg at week 15. In comparison, wildtype rats showed a visual acuity of 0.53 ± 0.01 cyc/deg and 0.49 ± 0.04 cyc/deg at week 7 and 15 respectively. We also determined that the visual acuity values determined by analysing the OMR using both eyes or single individual eye are comparable (Supplementary Figure 5B). Overall, our results highlight that P23H-3 rats experience a decrease in contrast sensitivity that can be detected through optomotor reflex tests starting from week 7 of age.

## Discussion

A variety of rodent models have been developed to replicate the human RHO-P23H mutation, the most common form of autosomal dominant RP. There has been prominent functional characterisation of P23H rodent models using ERGs, with a lesser focus on OMR characterisation. In the present study, we employed both ERG and OMR to characterise and compare over time the functional degeneration of the *Rho^P23H/WT^* mouse and rat models that have previously been identified to most closely resemble human pathophysiology. In scotopic conditions, our results demonstrated that P23H rodent models exhibited only minor decreases to OMRs despite ERG responses being substantially reduced. Similarly, OMR revealed visual acuity in photopic conditions were unchanged from wildtype in *Rho^P23H/WT^* mice, and even increased in P23H-3 rats, despite photopic ERGs being significantly reduced. These findings highlight that electrophysiological dysfunction via ERG testing precedes OMR disturbances in both mouse and rat *Rho^P23H/WT^*models.

### Disease modelling P23H mutations

Thorough characterisation of disease models is essential to ensure accurate modelling of human pathophysiology, and to provide a reliable platform for evaluating potential therapies.

Such analyses must consider that differences in animal background strains can significantly influence photoreceptor health. The most notable example of differences in background strain is that albino P23H-3 rats exhibit greater retinal degeneration than pigmented P23H-3 rats (Lowe et al., 2005, LaVail et al., 2018, McGill et al., 2012). In P23H-3 SD rats, by approximately 13 weeks of age, ONL thickness is already 60% that of wildtype (LaVail et al., 2018). By crossing those same P23H-3 SD rats with pigmented LE rats, ONL thickness is not largely distinguishable from wildtype at the same age (McGill et al., 2012). Similar to the latter study, we crossed P23H-3 SD x LE rats for optimal OMR testing and analysis. This progressive degeneration rate is exemplified by our findings that by 13 weeks the ONL thickness in P23H-3 rats has been significantly reduced to 82% of the ONL in wildtype rats. Similarly in mice, photoreceptor degeneration is exacerbated in albino mice with opsin mutations compared to pigmented mutant mice (Naash et al., 1996). There are retinal spatial distribution differences between albino and pigmented mice (Ortín-Martínez et al., 2014), with C57BL6/J having 70% more cone photoreceptors than albino mice (Whitney et al., 2011), which can have an effect on visual performance. Despite the prominent use of albino rodent models in medical research, in the care of vision research, these models possess a photoreceptor sensitivity that may not accurately represent the human condition as pigmented rodents. Consistent with this, our results highlight a low optomotor response in albino rats.

### ERG dysfunction precedes OMR decreases

The analysis of both mouse and rat P23H models, along with the rapid degeneration *rd1* mouse model, allowed us to monitor and compare visual performance with increasing degeneration severity. Although there were reductions in *Rho^P23H/WT^* mice scotopic ERG response at all ages tested, scotopic OMR was not significantly decreased until 2 months of age, when rods are reportedly half the number of wildtype (Sakami et al., 2011). These findings were mirrored in P23H-3 rats, as scotopic ERG responses were decreased but the scotopic OMR did not continue to decrease from 7 to 15 weeks of age. Persistent OMR in *Rho^P23H/WT^* mice has previously been hypothesised to be a result of potentiation of the rod-to-rod bipolar signal transmission as a compensation mechanism (Leinonen et al., 2020). Although this may be a potential factor contributing to persistent scotopic OMR *Rho^P23H/WT^*rodents, we have shown in *rd1* mice that in photopic conditions there is a residual OMR despite complete depletion of ERG responses at P24. Instead, we suggest that OMRs have a higher sensitivity to residual photoreceptor activity compared to full-field ERG.

Interestingly, our data shows photopic ERG a-waves of *Rho^P23H/WT^*mice were similar to wildtype, but b-waves were significantly reduced across the majority of ages tested. This reduction of the photopic b-wave in *Rho^P23H/WT^* mice has been presented in previous studies (Barwick et al., 2023, Vasireddy et al., 2011), whilst another study found no change in photopic b-wave by P70 (Wang et al., 2024). Photopic OMRs measuring visual acuity were unchanged across time and comparable to wildtype, reflecting that there were minimal adverse effects on cone-mediated responses during rod degeneration in *Rho^P23H/WT^* mice. In P23H-3 rats, our findings suggest that contrast sensitivity was significantly decreased starting at 7 weeks of age compared to wildtype, however visual acuity was not significantly reduced. Curiously, while visual acuity was slightly decreased in wildtype rats, in P23H-3 rats the visual acuity was slightly increased from week 7 to week 15. This result is different to previous findings possibly due to experimental differences, batch variations and/or variations in the OMR system used, which further highlighted the importance of independent validation studies. Indeed, a previous study by McGill et al. (2012) conducted OMR testing with slight variations in threshold sensitivity, therefore requiring data normalisation between the laboratory differences (McGill et al., 2012). In addition, their study used the OptoMotry system (CerebralMechanics), whilst here we used the OptoDrum (Striatech), likely contributing to experimental differences. Also using the OptoMotry system, Segura et al. (2018) showed a progressive decrease in VA in pigmented P23H-1 rats, which present a faster photoreceptor degeneration rate compared to the P23H-3 line used in our study (Segura et al., 2018).

Overall, this study provided a detailed charaterisation of the P23H rodent models and presented the first Optodrum OMR analysis of the P23H mouse and rats, with direct comparison to ERG data. Our findings have implications for the interpretation of visual function in preclinical models of retinal disease and highlight the need for behavioral analysis that extends beyond conventional ERG recordings. In particular, OMR testing can provide a useful endpoint for evaluating therapeutic interventions during late-stage disease progression where ERG responses may be unmeasurable.

## Methods

### Animals

The study was conducted in accordance with the ARVO Statement for the Use of Animals in Ophthalmic and Vision Research. All mouse lines were bred on a C57BL/6J background and were bred and housed at the Harry Perkins Institute of Medical Research bioresources facility. Animals were housed in a 12/12 h day/night cycle and received *ad libitum* access to food and water. Mouse experiments were approved by the Harry Perkins animal ethics committee and the University of Western Australia ethics committee (AE198). Homozygous P23H mice were obtained from The Jackson Laboratory (#017628) and used in a heterozygous form by crossing with C57BL/6J mice without mutation. The retinal degeneration 1 (*rd1*) mice were used as an autosomal recessive RP comparison, and the mutation was isolated from the Chrnb4.GFP line(Siegert et al., 2009) as part of a previous C3H/HeJ background. Pigmented heterozygous P23H rats were bred by crossing homozygous transgenic albino P23H line 3 rats with pigmented Long-Evans rats (Ozgene, Australia). All rats were bred and housed at EMSU (St. Vincent’s Hospital) animal facility and housed in a 12/12 h day/night cycle. Rat experiments were approved by the Animal Ethics Committee of St Vincent’s Hospital Melbourne (024/21).

### Electroretinogram (ERG)

To assess retinal function, full-field electroretinograms (ERGs) were recorded for scotopic (rod-mediated vision) and photopic (cone-mediated vision) responses using the Celeris ERG (Diagnosys, USA). Animals were dark adapted overnight and handled under dim red light for the scotopic paradigm. Animals were anaesthetised through intraperitoneal injection: 1) mice received 80 mg/kg ketamine (Ceva Animal Health Pty Ltd, Australia) and 10 mg/kg Xylazil-100 (Troy Laboratories, Australia); 2) rats were injected with 75 mg/kg ketamine and 10 mg/kg Xylazil-100. Animals received 1% tropicamide drops (Alcon, USA) to the eye to dilate the pupil, and rats received an additional drop of 2.5% phenylephrine hydrochloride (Bausch & Lomb, Canada). Following this, 2% hypromellose (HUB Pharmaceuticals LLC, USA) was applied to moisturise the eye and act as a contact fluid. Animals were placed on the Celeris ERG heated platform and electrodes were positioned in front of the eye, being careful to avoid contact with the cornea to reduce the occurrence of corneal scarring. Readings were measured using 1 millisecond flashes repeated four times and spaced 10 seconds apart. The scotopic light intensities measured for mice were 0.1, 1, 3, 10 and 25 cd.s/m^2^, with 60 seconds of recovery between different flash intensities. Following scotopic recordings, mice were light adapted for 10 minutes at 40 cd.s/m^2^ before commencing photopic testing. Photopic readings were measured at 1, 3, 5, 10 and 25 cd.s/m^2^ with a background light intensity of 30 cd.s/m^2^. Similar testing conditions were maintained for the rats, although the light intensities for scotopic recordings varied slightly, measuring 0.1, 0.3, 1, 3, 10 and 30 cd.s/m^2^. Oscillatory potentials (OPs) were generated for scotopic responses. Peak amplitudes of the first three oscillatory potentials were measured. Data was analysed for amplitudes of a-wave, b-wave, and OPs using the Espion V6 software (Diagnosys, USA) and Microsoft Excel.

### Optomotor response (OMR)

Optomotor responses (OMR) were assessed using the automated OptoDrum machine (Striatech, Germany), a closed box containing four LCD monitors that simulate a rotating cylinder of alternating white and black square-wave gratings. Animals were individually placed on a raised platform at the centre of the box, with a camera directly above to track temporal-nasal head movement (i.e OMR) via the computer-controlled software. The velocity of the virtual cylinder rotation was kept constant at 12°/s. Contrast sensitivity was determined under scotopic conditions, whereby the contrast between black and white stripes ranged from 99.97% to 0%, using the automated staircase algorithm of the OptoDrum software. In mice, contrast sensitivity was tested at a constant spatial frequency of 0.0556 cycles/degree, whilst rats were tested at a constant spatial frequency of 0.083 cycles/degree. Visual acuity was analysed under photopic conditions, with the contrast between stripes kept constant at 99%. The spatial frequency threshold was automatically determined using the staircase algorithm from the OptoDrum software, ranging from 0.0556 to 0.556 cycles/degree. Animals were tested in each condition until there was an absence of a head movement reflex in response to the visual stimulus. This was determined by the software if the head movement score no longer exceeded the chance level of a stimulus-independent head movement.

### Optical coherence tomography (OCT)

The structure of the retina was analysed in P23H-3 rats via optical coherence tomography using a Spectralis HRA+OCT machine (Heidelberg Engineering GmbH, Germany). High resolution horizontal volume scans were taken across the retina with the optic nerve head at the centre. Each scan was a composite image average of 23 frames to correct projection artefacts. The OCT images were analysed using the segmentation tool from the Spectralis software. Retinal thickness values are the average of 4-8 areas from the retinal thickness map, avoiding the optic nerve head. The outer nuclear layer (ONL) thickness values represent the measurements of the manually annotated ONL layer marker, the region from the outer plexiform layer to the external limiting membrane.

### Statistical analysis

Data was analysed using the GraphPad Prism 9 software and presented as mean ± standard error of the mean (SEM). Mouse data that consisted of more than two groups was analysed using a two-way ANOVA and Tukey’s multiple comparisons post-hoc test. Rat data was analysed using a one-way ANOVA. Data was considered significant if *P* < 0.05.

## Acknowledgements

This work is supported by the University of Melbourne Early Career Researcher grant (DUC). RCBW is supported by the University of Melbourne, the Centre for Eye Research Australia and the Medical Research Future Fund (MRF2024365). The Centre for Eye Research Australia receives operational infrastructure support from the Victorian Government. The authors acknowledge the facilities, and the scientific and technical assistance of the Australian Microscopy & Microanalysis Research Facility at the Centre for Microscopy, Characterisation & Analysis, The University of Western Australia, a facility funded by the university, State and Commonwealth Governments.

## Author contributions statement

Study design and concept: AAB, DUC, ARH, RCBW, LSC; Experiment: AAB, DUC, GH, SC, REJ; Analysis: AAB, LW, DUC, RCBW; Funding: RCBW, LSC; Manuscript writing: AAB, DUC, REJ. All authors approved the final manuscript.

## Supplementary Materials

**Supplementary Figure 1.**
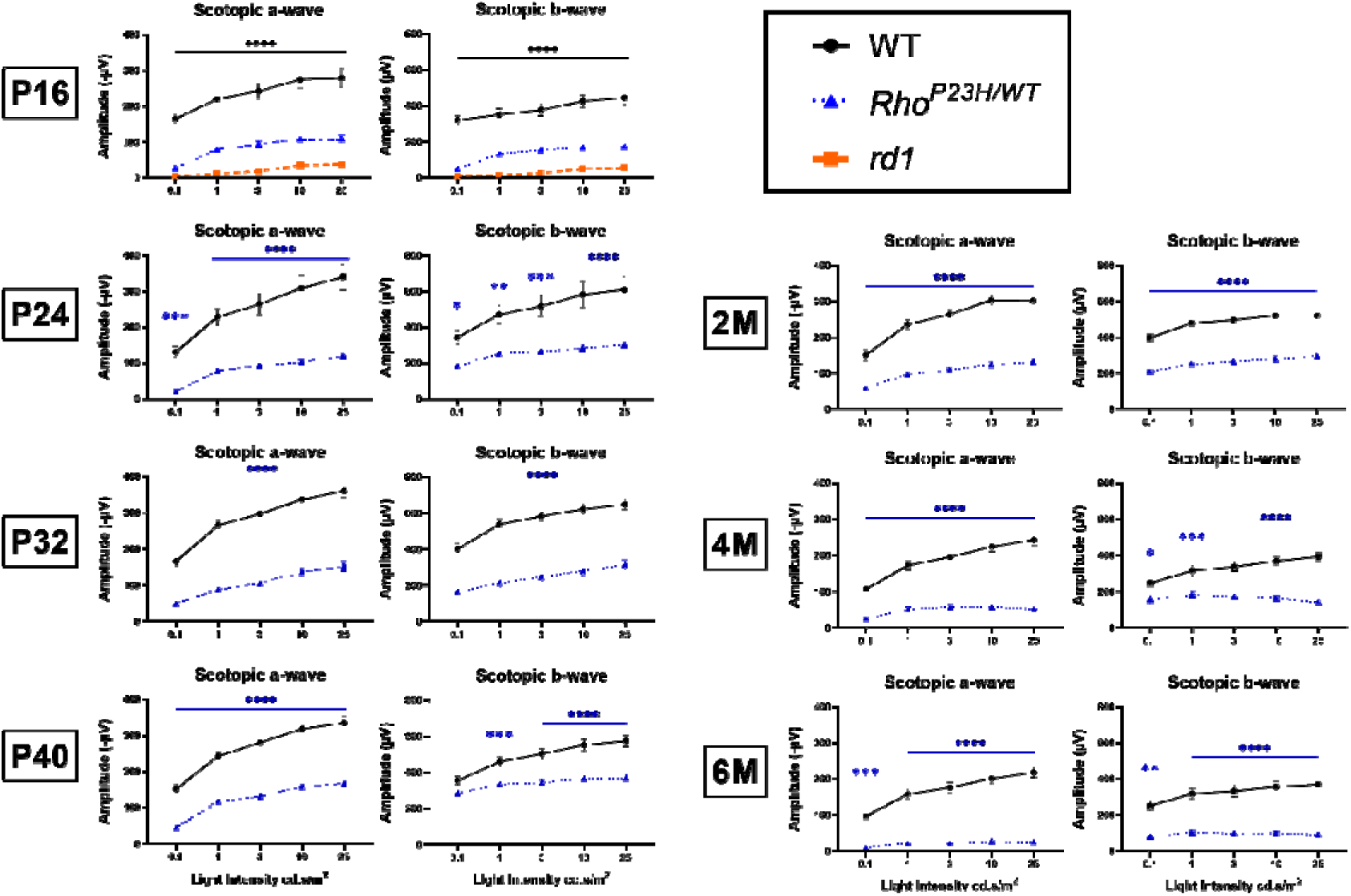
Scotopic ERG responses at all light intensities across age groups in mice. Black asterisks indicate both *Rho^P23H/WT^* and *rd1* ERGs are significantly decreased compared to wildtype, whilst blue asterisks indicate significant decreases in *Rho^P23H/WT^* to wildtype. *Rd1* mice no longer had a measurable scotopic ERG response after P16 and are therefore not shown. Numbers are mean ± SEM. *n* = 3-6 per age per line. \**P* < 0.05, \*\**P* < 0.01, \*\*\**P* < 0.001, \*\*\*\**P* < 0.0001, twoway ANOVA and Tukey’s multiple comparisons test when comparing between all 3 lines, and Sidak’s multiple comparisons test when comparing just *Rho^P23H/WT^*and wildtype. 2M: 2 months; 4M: 4 months; 6M: 6 months

**Supplementary Figure 2.**
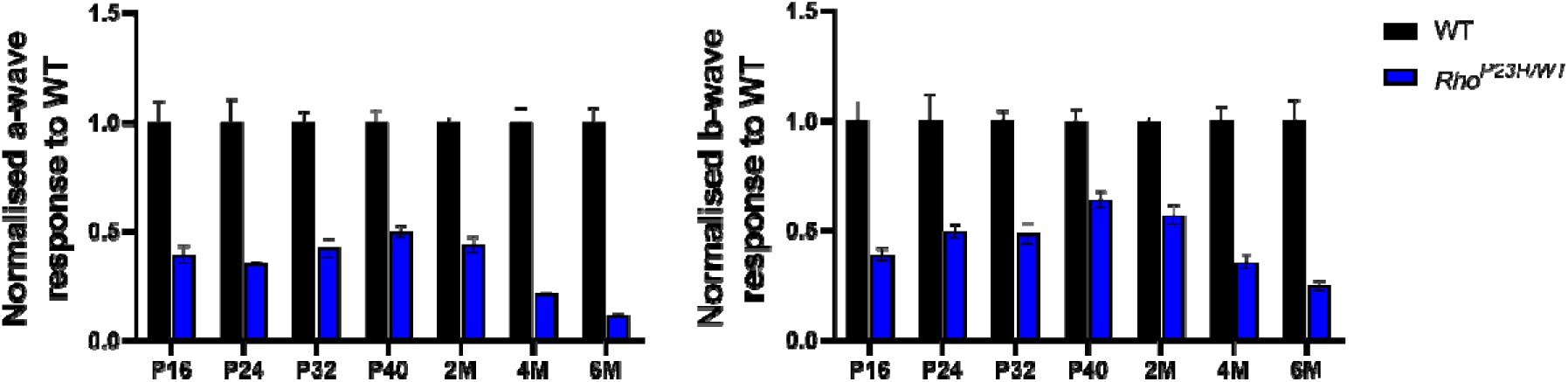
Scotopic ERG responses in *Rho^P23H/WT^* mice normalised to wildtype. (**A**) scotopic a-wave and (**B**) b-wave responses of *Rho^P23H/WT^* mice (blue) normalised to wildtype mice (black) at 25 cd.s/m^2^ light intensity across ages. Numbers are mean ± SEM. *n* = 3-6 per age per line. 2M: 2 months; 4M: 4 months; 6M: 6 months.

**Supplementary Figure 3.**
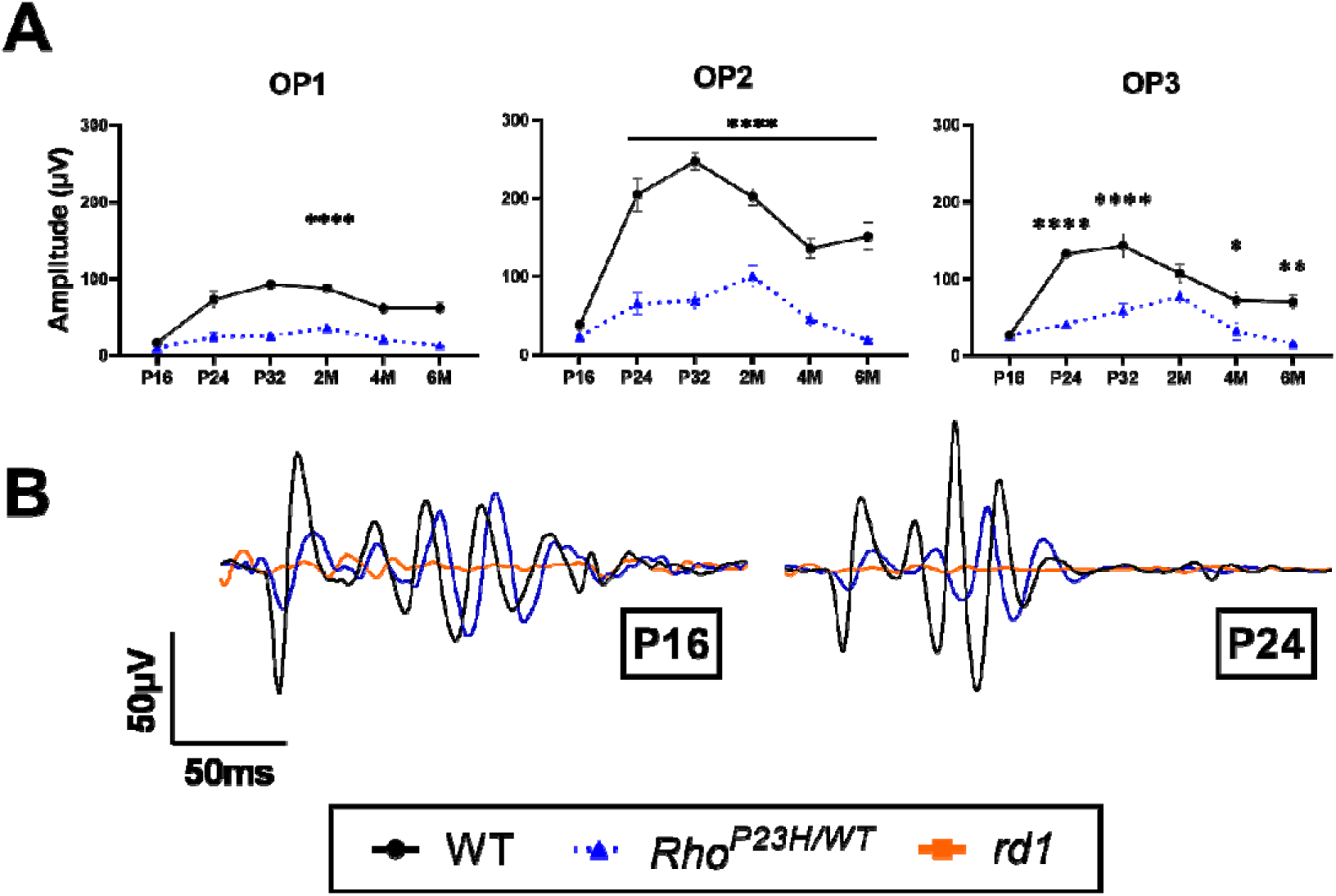
Scotopic OP responses across age groups. (**A**) OP1, OP2, and OP3 responses in *Rho^P23H/WT^* mice compared to wildtype. *Rd1* OPs were excluded as they were not quantifiable at all ages. (**B**) Representative traces of scotopic OP responses are shown for each line at P16 and P24. Numbers are mean ± SEM. *n* = 3-6 per age per line. \**P* < 0.05, \*\**P* < 0.01, \*\*\*\**P* < 0.0001, two-way ANOVA and Sidak’s multiple comparisons test. 2M: 2 months; 4M: 4 months; 6M: 6 months.

**Supplementary Figure 4.**
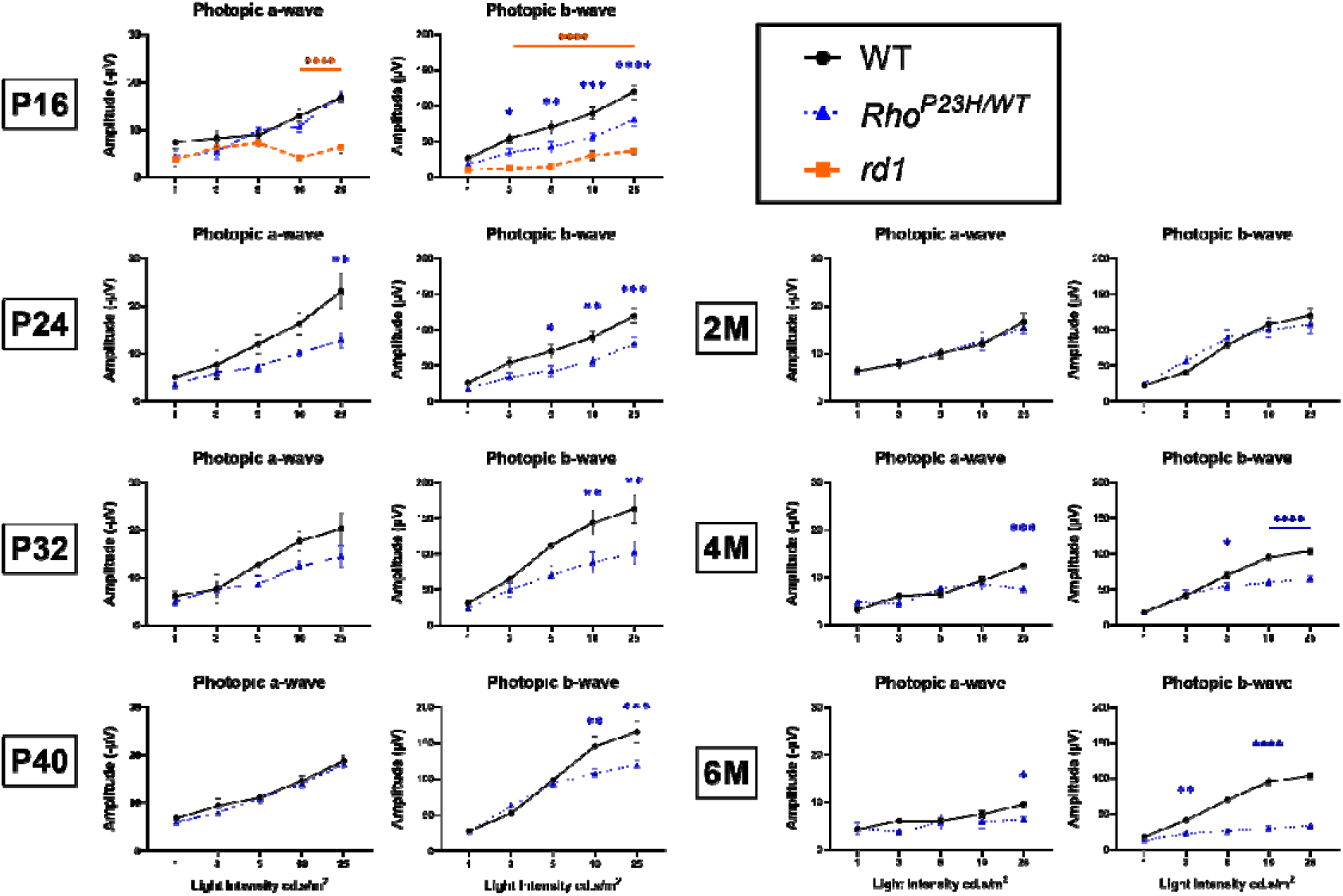
Photopic ERG responses at all light intensities across age groups. Blue asterisks indicate significant decreases in *Rho^P23H/WT^* to wildtype, and orange asterisks indicate significant decreases in *rd1* to wildtype. *Rd1* mice no longer had a measurable photopic ERG response after P16 and are therefore not shown. Numbers are mean ± SEM. *n* = 3-6 per age per line. \**P* < 0.05, \*\**P* < 0.01, \*\*\**P* < 0.001, \*\*\*\**P* < 0.0001, two-way ANOVA and Tukey’s multiple comparisons test when comparing between all 3 lines, and Sidak’s multiple comparisons test when comparing just *Rho^P23H/WT^* and wildtype. 2M: 2 months; 4M: 4 months; 6M: 6 months.

**Supplementary Figure 5.**
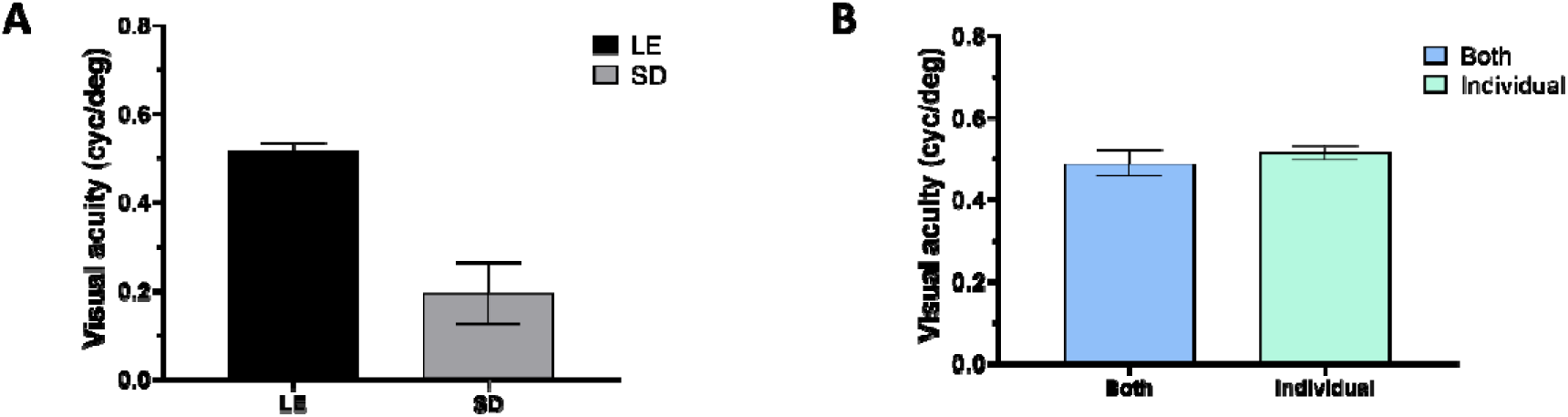
Visual acuity test optimization in rats. (**A**) Visual acuity test in pigmented (LE) and albino (SD) wild type rats. (**B**) Comparison of the visual acuity thresholds determined by testing both eyes and each eye as individual subjects. Numbers are mean ± SEM. *n* = 8-16 eyes.

**Supplementary Figure 6.**
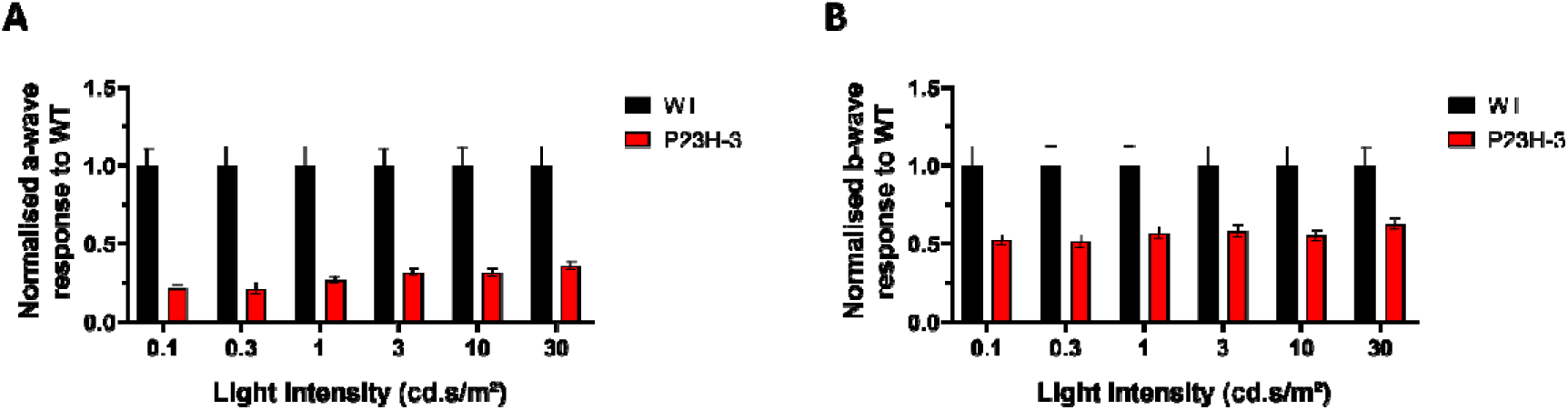
Normalised scotopic ERG response in 13 week old P23H rats. (**A**) scotopic a-wave and (**B**) b-wave responses of P23H-3 rats (red) normalised to the response of WT rats (black) at flashes of 0.1, 0.3, 1, 3, 10 and 30 cd.s/m^2^. Numbers are mean ± SEM. *n* = 10-12 eyes.

